# Robust and Brain-Like Working Memory through Short-Term Synaptic Plasticity

**DOI:** 10.1101/2022.01.09.475558

**Authors:** Leo Kozachkov, John Tauber, Mikael Lundqvist, Scott L. Brincat, Jean-Jacques Slotine, Earl K. Miller

## Abstract

Working memory has long been thought to arise from sustained spiking/attractor dynamics. However, recent work has suggested that short-term synaptic plasticity (STSP) may help maintain attractor states over gaps in time with little or no spiking. To determine if STSP endows additional functional advantages, we trained artificial recurrent neural networks (RNNs) with and without STSP to perform an object working memory task. We found that RNNs with and without STSP were able to maintain memories over distractors presented in the middle of the memory delay. However, RNNs with STSP showed activity that was similar to that seen in the cortex of a non-human primate (NHP) performing the same task. By contrast, RNNs without STSP showed activity that was less brain-like. Further, RNNs with STSP were more robust to noise and network degradation than RNNs without STSP. These results show that STSP can not only help maintain working memories, it also makes neural networks more robust and brain-like.

## Introduction

Working memory (WM), the holding of information “online” and available for processing, is central to higher cognitive functions^1,2^. Its well-established neural correlate is spiking over a memory delay^3–5^. For many years, this was thought to be the sole mechanism underlying WM maintenance. The idea is that sensory inputs elicit unique patterns of spiking that are sustained via recurrent connections^6^, creating attractor states—stable patterns of activity that retain the WM^7^. It seems evident that these attractor dynamics play an important role in WM. Recent observations, however, have suggested that there may be more going on ^8–11^. A few neurons seem to show spiking that looks persistent enough to be an attractor state, but the bulk of neurons show memory delay spiking that is sparse^12–15^. This is especially true when spiking is examined in real time (i.e., on single trials) because averaging across trials can create the appearance of persistence even when the underlying activity is quite sparse. Alternatively, it might just be that attractor dynamics are more a phenomenon of large populations of neurons rather than single neurons. By this we mean that attractors may be easy to spot when we have access to all the neurons, but hard to spot when we only subsample a tiny fraction.

This all begs the question of how WMs are maintained over these gaps in time with little-to-no spiking^11,16^. One possibility was suggested by observations of short-term synaptic plasticity (STSP) in circuits in the prefrontal cortex, transient (< 1 second) changes in synaptic weights induced by spiking^17^. Several groups have suggested updating the attractor dynamics model with this feature^18–20^. The idea is that STSP helps the spiking. Spikes induce a transient “impression” in the synaptic weights that can maintain the network state between spikes ^21–23^. Evidence for STSP comes from techniques like patch-clamp recording that are difficult to implement in the working brain, especially in NHPs. Thus, we tested the role of STSP in WM by using computational modeling in conjunction with “ground-truthing” via analysis of spiking recorded from the PFC of a NHP performing a WM task. We aimed to determine if network models with STSP can solve the working memory task, whether they have properties similar to those seen in the actual brain, and whether STSP endows functional advantages. The answer to these questions was “yes”.

We trained Recurrent Neural Networks (RNNs) with and without STSP to test how it affects network performance and function. We focused on the key property of robustness^9,24–27^. Working memories must be maintained in the face of distractions. Networks need to deal with noise and show graceful degradation (i.e., continue to function when portions of the network are damaged). Our analysis showed RNNs with and without STSP were robust against distractors. However, *only the RNNs with STSP were “brain-like”*—their activity more closely resembled activity recorded from the prefrontal cortex of a NHP performing a WM task. RNNs with STSP were also more robust against increasing levels of noise and synaptic ablation. Thus, STSP explains how WM can be maintained between spiking. It also carries additional functional advantages, such as robustness to various perturbations.

## Results

A NHP was trained to perform an object delayed-match-to-sample task (Figure 1). The NHP was shown a sample object and had to choose its match after a variable-length memory delay. At mid-delay a distractor object (1 of 2 possible objects never used as samples) was presented (for 0.25s) on 50% randomly chosen trials. We recorded multi-unit activity (MUA) bilaterally in dorsolateral PFC (dlPFC) and ventrolateral PFC (vlPFC) using four 64-electrode Utah arrays for a total of 256 electrodes. The animal learned to do the task consistently at ∼99% accuracy for both distractor and non-distractor trials

**Figure 1:**
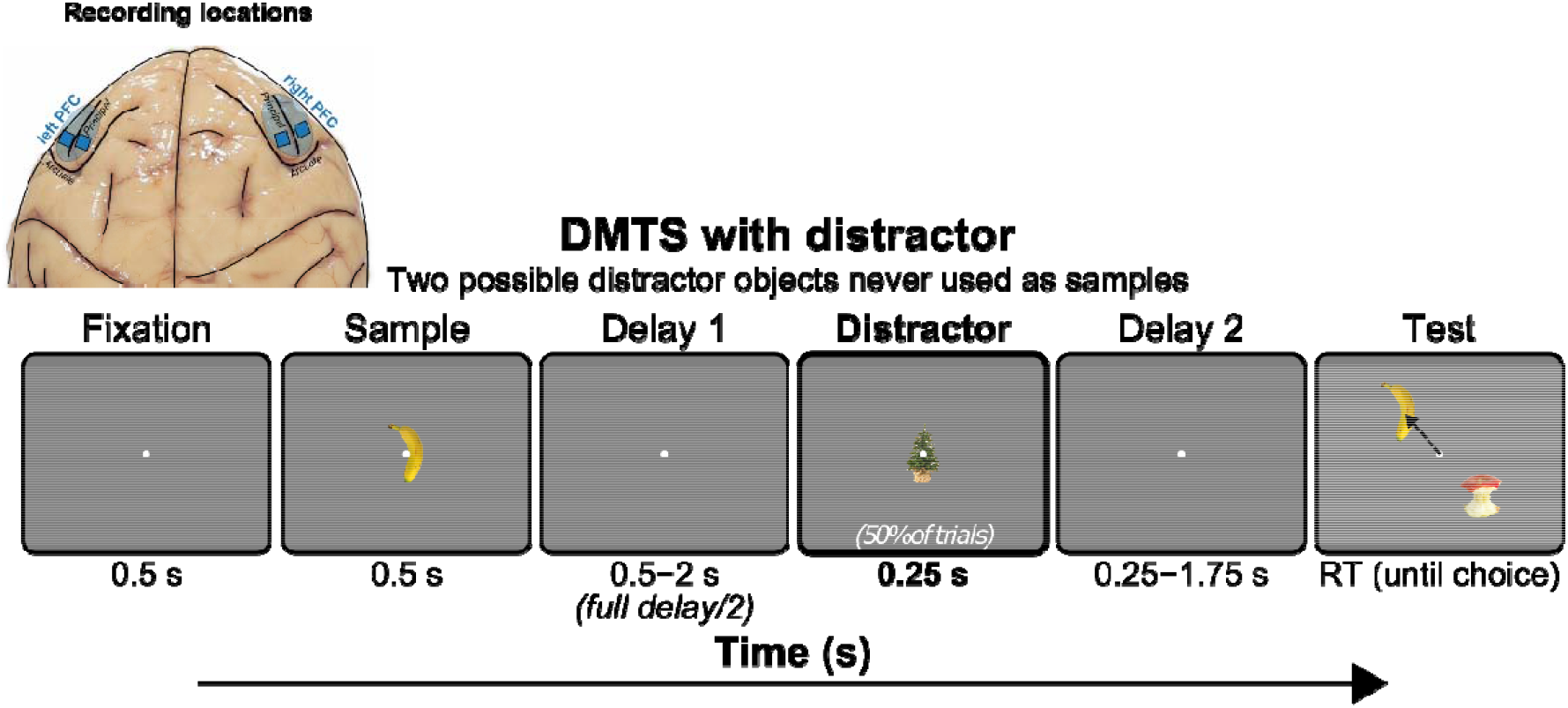
Electrode location and task structure. Utah arrays were implanted bilaterally in dorsolateral PFC (dlPFC) and ventrolateral PFC (vlPFC). Animal performed a distracted delayed match-to-sample task. Each trial began with visual fixation on the middle of the screen for 0.5s. Fixation was maintained throughout the trial until the behavioral response. The delay length was parametrically varied from 1 – 4 s in five logarithmic steps, randomly chosen each trial. At mid-delay a neutral distractor (1 of 2 possible objects never used as samples) was presented randomly on 50% of trials. During the multi-choice test the NHP was allowed to freely saccade between all objects on the screen. The final choice was indicated by fixating on it for at least one second.

### Sample Information in Population Neural Activity Was Weak Over Longer Delays

First, we examined MUA recorded from the lateral PFC. To quantify the amount of sample object information carried by spiking, we used a linear classifier (see methods for details)^1^. This showed that spiking carried sample object information for about one second after the sample disappeared. From the start of the delay period the decoder accuracy decreased steadily towards chance (Figure 2a, Figure S 1). We corroborated this by measuring the distance between neural population activity for all pairs of sample objects. That gave an average distance between experimental conditions at every timepoint (see methods). This showed that the distance between population MUA activity for different samples returned to pre-stimulus levels (Figure 2b, Figure S 2). Interestingly, we found that this was *not* simply due to spiking returning to pre-sample values. We determined this by training a classifier to discriminate between pre and post sample spiking activity. We found that this classifier was consistently able to discriminate between pre and post sample spiking activity over the delay (Figure S 3).

**Figure 2:**
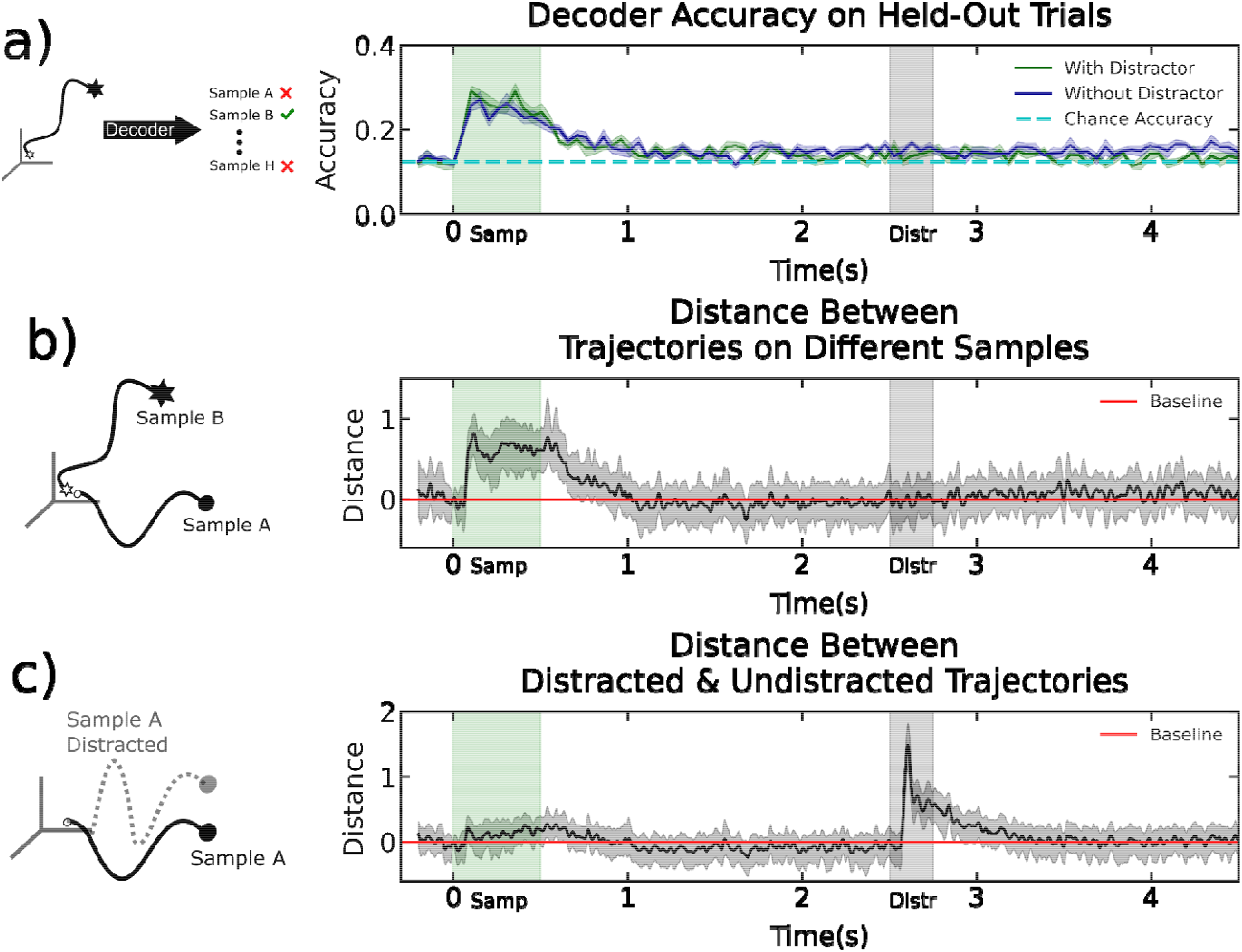
a) Left: training a decoder to predict sample identity given a neural trajectory. Right: The decoder accuracy on held-out trials for distracted vs. undistracted trials b) Left: comparing trial-averaged trajectories corresponding to different samples. Right: The average pairwise distance in state-space between trajectories elicited by all possible sample images. Normalized by the average pre-stim distance. c) Left: comparing trial-averaged distracted vs. non-distracted trajectories through neural state space. Right: The distance in state space between distracted vs. non-distracted trajectories throughout the trial. Shown are trials with a delay of four seconds.

### Population Neural Activity Was Robust to Distractors

We found that when the mid-delay distractor was presented, neural trajectories diverged from that of non-distractor trials (Figure 2c, Figure S 4). Once the distractor disappeared, the trajectories quickly reconvened, indicating neural stability. This was true for all five delay lengths used (1 – 4 s in five logarithmic steps, Figure S 4). The time course of trajectory reconvening was roughly exponential with a time constant of ∼200 milliseconds (Figure 2c, Figure S 4). We determined this by fitting an exponential function to the recovery curves and measuring the inverse of the fitted decay constants. This is consistent with prior observations that time constants in cortex peak at this value in the PFC ^28^.

This all raises a couple of questions. 1. How can PFC networks support WM task performance when, across the population, sample information in spiking is relatively weak? 2. How do PFC networks achieve the stability to recover from distractors? To answer these questions, we used RNN modeling and neural network theory. As we will show, the two questions share a common answer: STSP.

### RNNs With STSP are More Brain-Like

RNNs with and without STSP were able to successfully perform the object delayed match to sample task. However, only the RNNs with STSP did so with activity that was similar to that seen in the actual PFC. Our hypothesis space consisted of four different kinds of RNNs: two with fixed synaptic weights (fixed after training) and two with STSP. The two fixed-weight networks were ‘vanilla’ RNNs. They differed only in their activation functions. One used a hyperbolic tan (tanh), the other used a rectified linear (ReLU). We will refer to these fixed synapse networks as *FS-tanh* and *FS-relu* respectively. We choose these two activation functions because they represent two sides of a spectrum. Tanh units can become unresponsive for very large inputs (i.e saturate) while ReLU units cannot. Both activations are commonly used throughout computational neuroscience and machine learning^29,30^. They lead RNNs to prefer one strategy over another for performing a task. ReLU units are ideal for forming line attractors, while tanh units are ideal for forming point attractors^30^.

The synaptic weights for the fixed-synapse networks are only adjusted during training. After training, they are untouched. The other two models use STSP to adjust synaptic weights during each trial. This represents the distinction between long-term and short-term memory. Long-term memory is acquired over the course of task-optimization and remains fixed throughout the trial. Short-term memory changes during the trial and reflects the contents of working memory. We reset the state of the plastic synapses at the beginning of each trial.

One RNN with STSP was based on a model introduced by Mongillo et al^19^. The model uses a set of simplified equations for synaptic calcium dynamics to endow the RNN with STSP. The Mongillo model adjusts synaptic weights based on presynaptic neural activity. For this reason we will call it *PS-pre*. The other RNN with STSP was introduced by Kozachkov et al.^31^ and adjusts synaptic weights in a way that depends on both the pre and post synaptic neural activity. For this reason we will call it *PS-hebb*. It used excitatory anti-Hebbian and inhibitory Hebbian mechanisms to stabilize the RNN^31^.

All models were trained with backpropagation-through-time using a standard deep learning library^32^. While the FS models were trained without any explicit constraints, the PS models needed to be parameterized to ensure they satisfy certain properties throughout training. In particular, the *PS-pre* model had separate excitatory and inhibitory neurons. The weight matrix therefore needed to be parameterized so that these populations always remained separate (see Methods). Likewise, for *PS-hebb* the weights were parameterized so the network remained stable throughout training. To ensure that our results did not depend on the particular choice of training hyperparameters, we trained the four models under a wide range of hyperparameter settings. These hyperparameters were: the degree of activity regularization and parameter regularization during training, the size of the network, and the decay rate of the synapses in *PS-hebb*. Roughly 2000 models were trained in total. We refer the reader to the Methods section for more details on the models and the training process.As in the analysis of the experimental data, we used a decoder to read out sample object information over time. We did this for both the fixed-synapse and plastic-synapse models. For the fixed-synapse RNNs we trained the decoder on trajectories from the spike rates. For the STSP RNNs, we trained two separate decoders: one on the spike-rates and one on the synaptic weights.

In all models, sample presentation elicited a large increase in decoding accuracy (Figure 3). Spike rate decoding of sample information in the fixed-synapse models remained high throughout the memory delay. This is in sharp contrast to the actual PFC MUA data. The PFC MUA showed a large drop in sample object decoding accuracy once the sample disappeared and especially for delays longer than one second. This was mirrored in the STSP models (Figure 3). However, sample decoding accuracy on the synaptic weights for the STSP models remained high over the entire delay, including longer delays (Figure 3).

**Figure 3:**
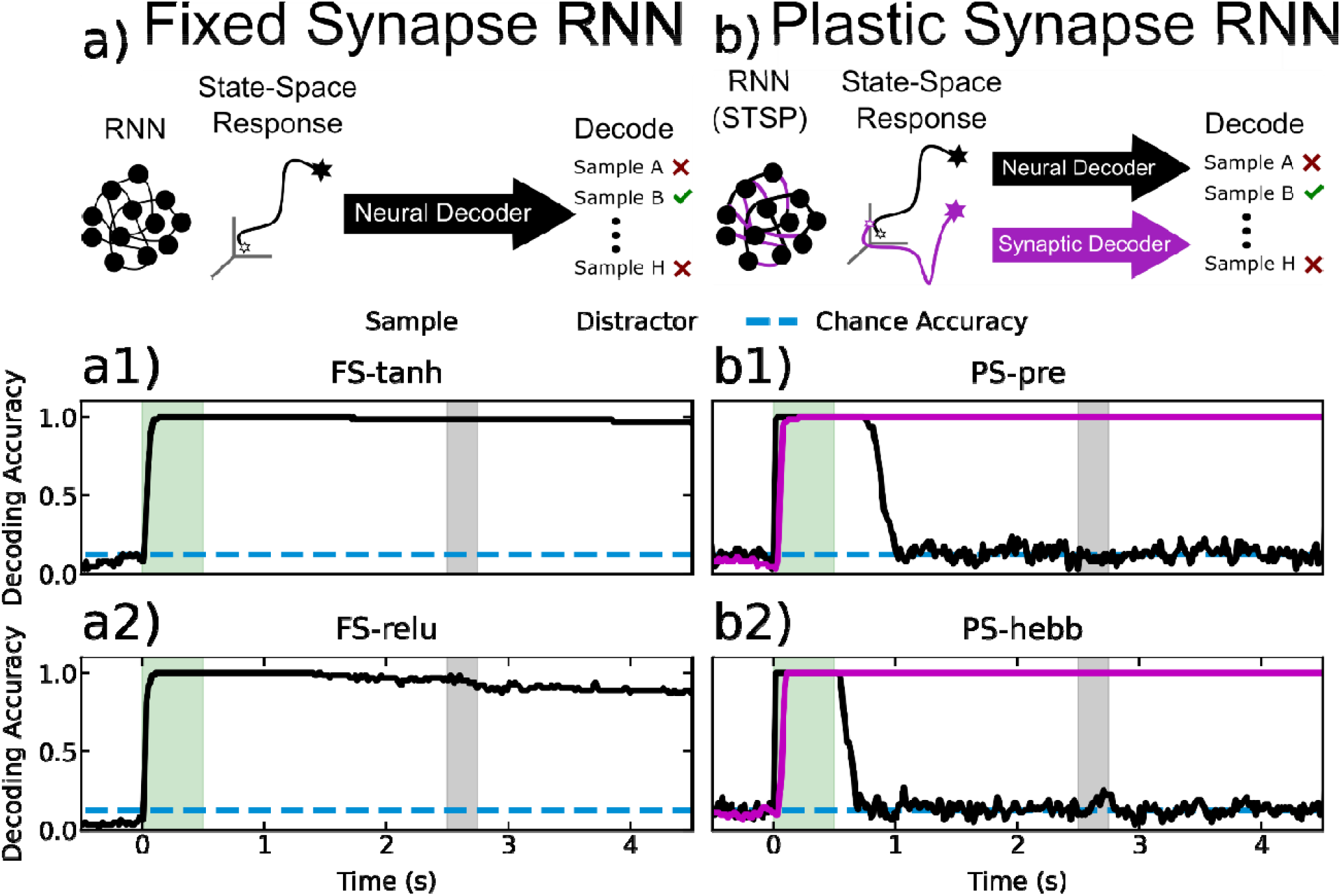
Decoding accuracy for RNNs. a) A decoder is trained to predict the sample label from RNN neural trajectories, for the fixed-synapse RNNs. a1) The accuracy of the trained decoder as a function of time from sample onset for FS-tanh. a2) The same plot as (a1), for FS-relu. b) Two decoders are trained on the neural and synaptic trajectories separately. Black lines indicate neural decoding, purple indicate synaptic decoding. b1) Neural and synaptic decoding accuracy as a function of time for PS-pre. b2) Same plot as in (b1) but for PS-hebb.

This raises the question: do STSP models better capture the dynamics of the real PFC? We addressed this question by comparing the similarity of each of the ∼2000 trained models to the actual neural data. To quantify the similarity between model and brain, we used the Pearson correlation between the decoder accuracy curves for the real neural data and the RNNs (Figure 4). We did this for each delay length and averaged the correlation scores to get a final brain-RNN correlation value for each model.

**Figure 4:**
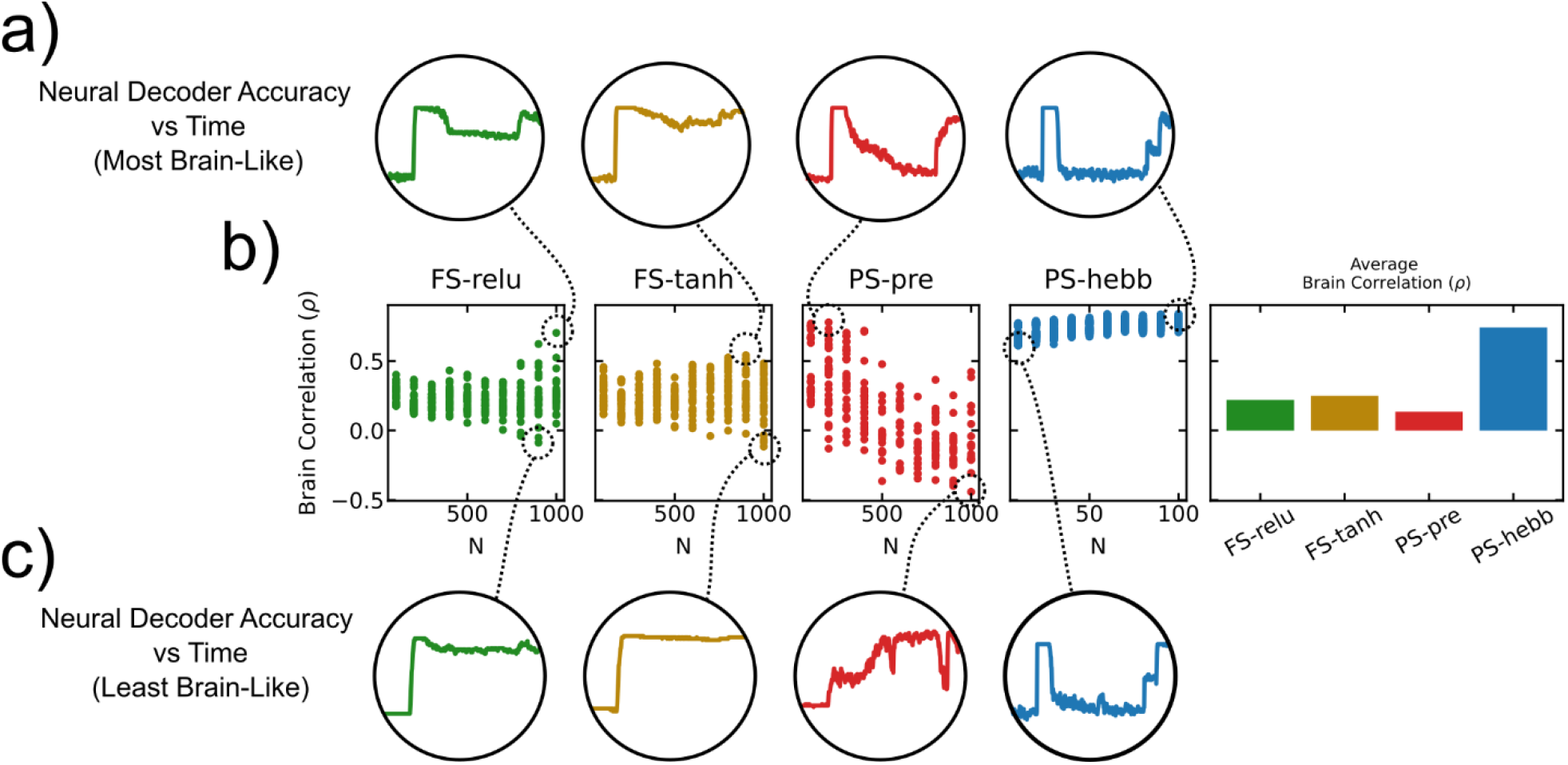
Different training hyperparameters lead to different levels of brain-similarity. Brain-similarity was measured as the correlation between the neural decoding accuracy curves for the real neural data and the RNNs. a) Neural decoder accuracy traces corresponding to the most brain-like RNNs. b) Three left-most subplots show the brain-similarity as a function of number of hidden units. Each dot represents an RNN trained with different hyperparameter settings. The rightmost subplot shows the average of all the brain correlation values. c) Same plot as (a), for the least brain-like RNNs.

This analysis revealed that the *PS-hebb* model was the most similar to the brain across a wide range of hyperparameters (Figure 4). Interestingly, this analysis also revealed that the *PS-pre* model ‘switched’ WM strategies as the number of neurons increased (Figure 4). The smaller networks that correlated highly with brain data had a low amount of persistent neural activity during the delay period. This indicates that information was primarily stored in the synaptic trace. The larger networks were less correlated with the brain data, and exhibited higher amounts of persistent neural activity (Figure 4). Within the fixed-synapse model class, the models that were most brain-like showed a marked dip in neural decoder accuracy during the delay period—as expected. However, this dip was far less severe than in the models with plastic synapses. The result was that models without synaptic plasticity had a lower similarity to the actual neural data.

An important difference between measuring information in RNNs and brains is that we subsample neurons in the brain. It is possible that the decoding accuracy curves are affected by this subsampling. To control for this, we repeated the above analysis while subsampling at four different values. We randomly picked 0.01%,1.0%,10.0% and 100.0% of the neurons in each RNN to decode from. The result was that STSP networks were more brain-like and this did not change because of subsampling (Figure S 5).

### STSP Increases Structural Robustness

The two models with STSP were *structurally* more robust than the models with fixed synaptic weights. To measure robustness, we examined how the performance accuracy of the trained network varied as we randomly ablated a varying fraction of synaptic weights (Figure 5). This showed that the STSP network’s performance on the task remained high even when as many as half of the synapses were ablated. By contrast, the fixed-synapse models were highly sensitive to synaptic ablation. Their performance severely degraded even with ablation of only 10-20% of the synapses.

**Figure 5:**
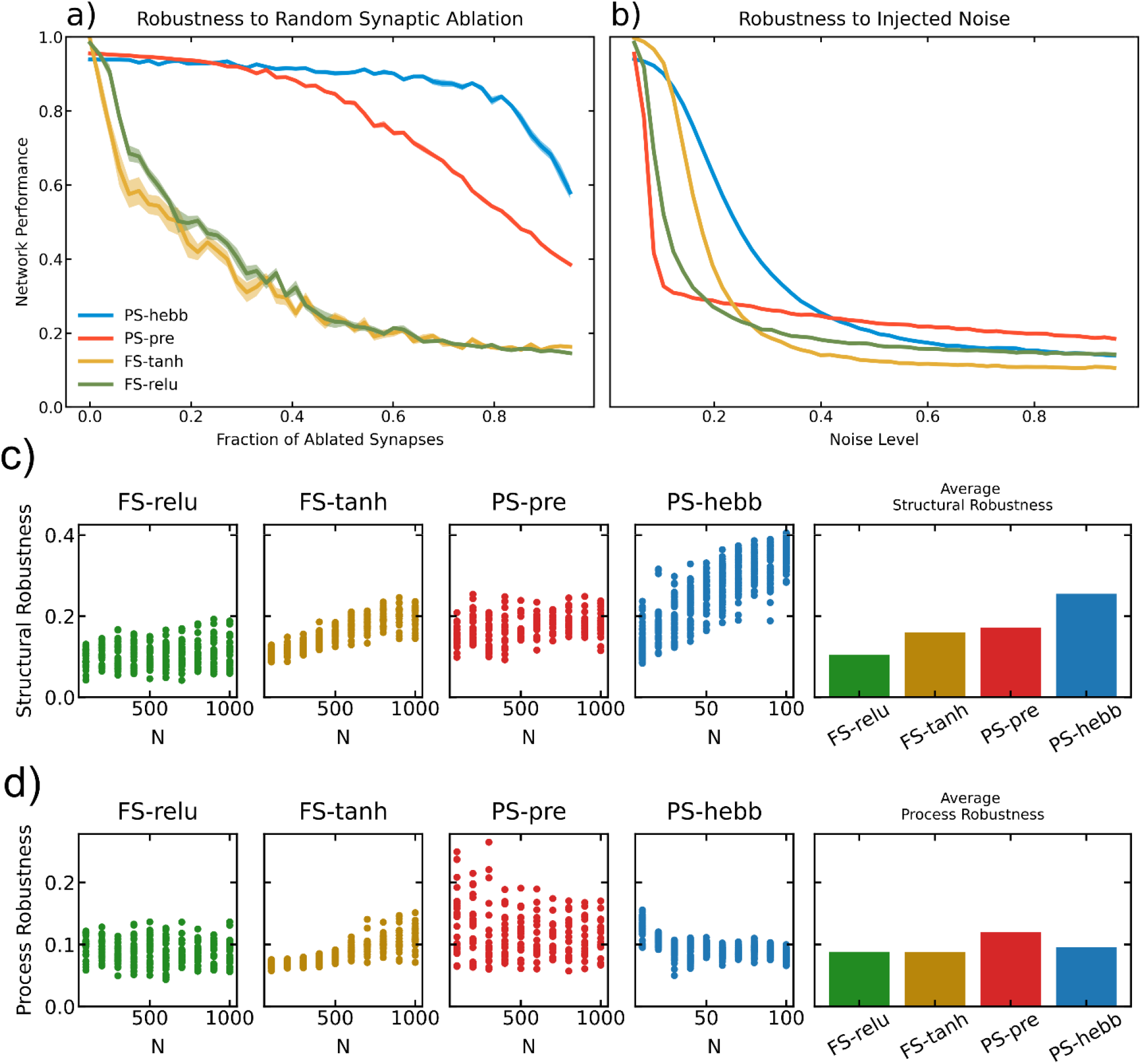
Measuring the robustness of the models. a) RNN test set performance as function of the fraction of randomly ablated synapses. Shaded regions denote standard error computed over 10 repetitions. b) Same as in (a), but as a function of increasing process noise. c) Structural robustness measured across all the models. For each model trained with different hyperparameter settings, a single robustness score was computed using the technique described in the main text. Right most subplot shows the average across all values. d) Same plot as in (c), but for process noise.

We repeated this analysis over all the models trained with different hyperparameters. From each model we defined two measurements to quantify robustness. The first was robustness to structural noise. The second was the robustness to synaptic noise. Both measurements were calculated in a similar way. We took the weighted average RNN performance over all the noise values tested. We weighed each term in the sum by the corresponding noise value. The reason for this weighting scheme is simple. If the RNN performance is high when the noise is low, this does not tell us much about robustness. However, if the RNN performance is high when the noise is high, this does indicate robustness. Thus, a natural measure of robustness is the product of noise and performance. This analysis revealed that the *PS-hebb* model was more robust to structural perturbations across a wide range of hyperparameters (Figure 5). This also revealed that *PS-pre* was most robust to process noise. To better understand how the different trained models achieved robust WM, we investigated how these networks organized their activity in state-space.

We found that the fixed-synapse RNNs learned to perform the task by using simple attractors. To visualize these attractors, we projected the high-dimensional RNN activity into a lower dimensional space using Linear Discriminant Analysis (Figure 6). We reasoned that this projection would give us the clearest visualization of the underlying state space attractors. For the plastic-synapse RNNs, we did this for both the neural and synaptic state-space. To better quantify the attractor properties of these networks, we measured the distances between state-space trajectories of the most brain-like models, as we did for the neural data (Figure 7). This revealed different trajectories for different sample objects. Each trajectory settled into a steady state that was unique for each sample, indicating the presence of attractors (Figure 6, Figure 7). Comparison between trials with and without a distractor showed that the distractor temporarily knocked trajectories out of the attractor state. Then trajectories quickly returned to the pre-distractor attractor state. This is how the fixed-synapse RNNs achieve robustness. Once the neural trajectories were ‘captured’ by the appropriate attractor, they remained fixed there and can return there after being perturbed. In terms of neural activity this means that the spike rates stayed at a particular, elevated firing rate across the delay period corresponding to the sample identity. In other words, these models achieve robust WM through persistent neural activity as a consequence of attractor dynamics, as observed in many prior models.

**Figure 6:**
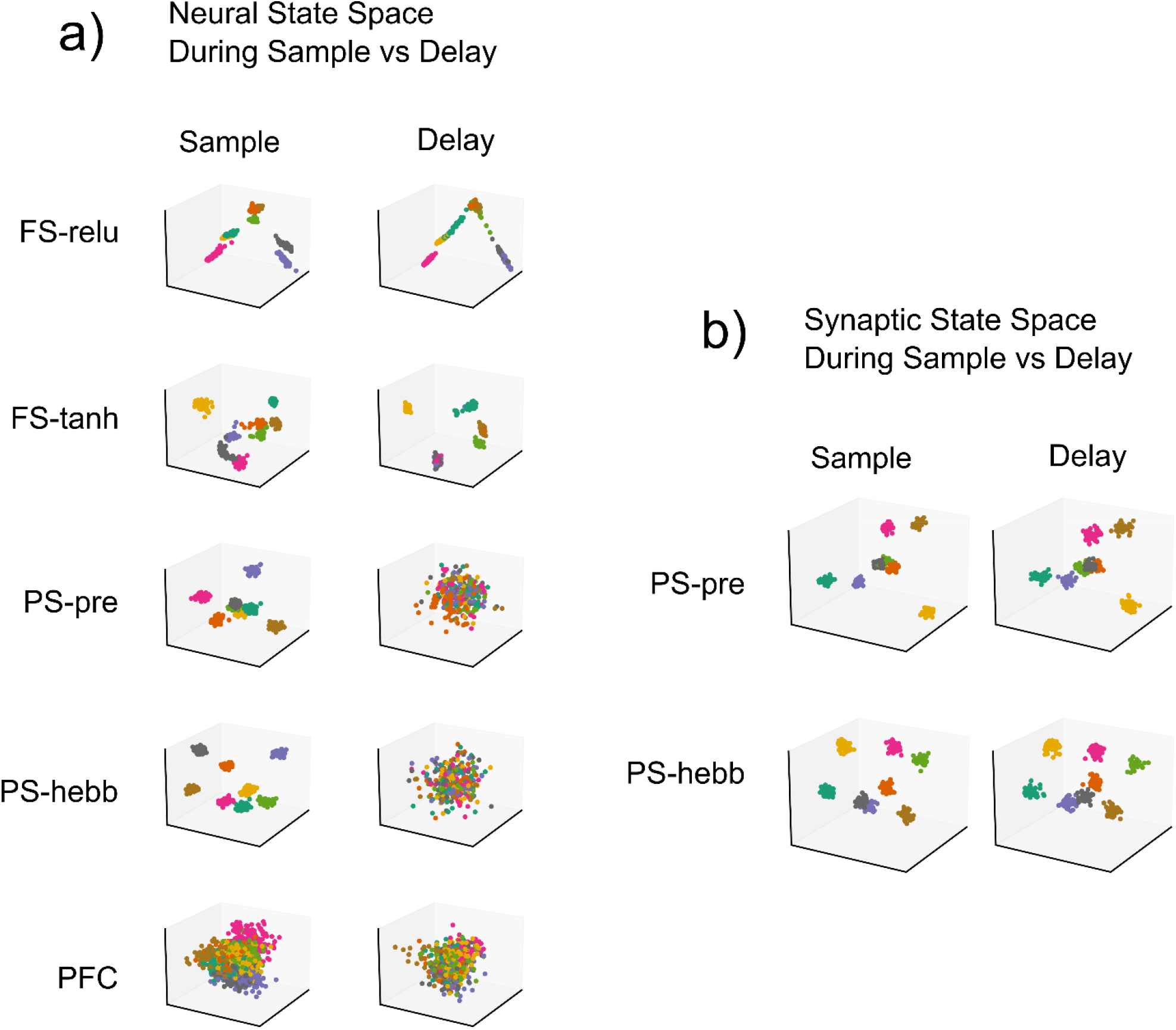
Dimensionality reduced space plots for the most brain-like models. We time-averaged RNN activity during the sample-period as well as 500ms before the end of the delay period. We used Linear Discriminant Analysis (LDA) to project the data into three-dimensions. To account for differences in training/test splits, we used an Orthogonal Procrustes operation to rotationally align the sample and delay period activity. Colors denote sample IDs. Fixed synapse models have activity organized around simple attractors in state space. There is one attractor for each sample ID. Plastic synapse models exhibit high sample-separability in synaptic state space, and limited separability in the neural state space during the delay period. Similarly, PFC exhibited higher neural discriminability during the sample than during the delay period.

**Figure 7:**
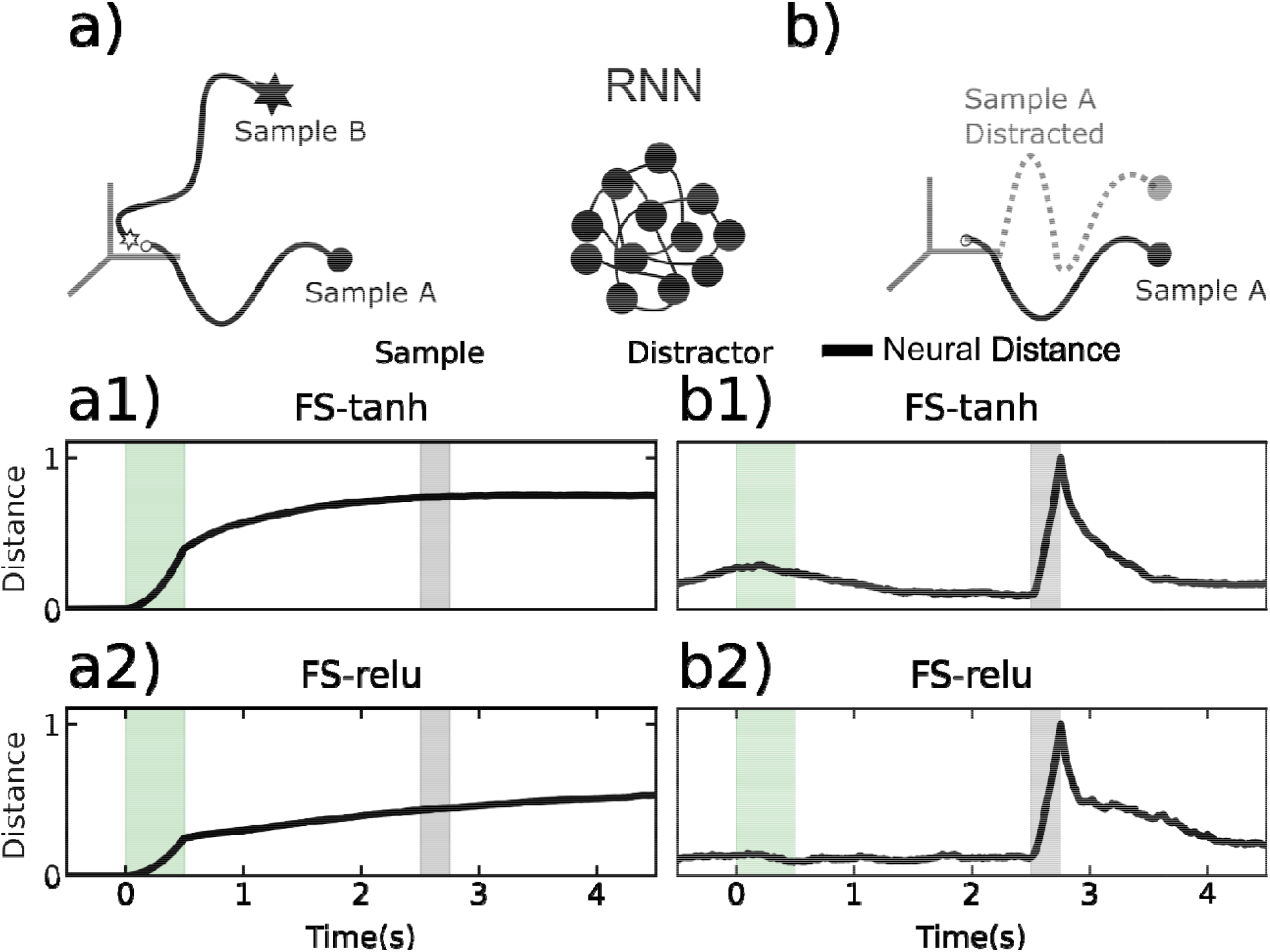
Distances between neural trajectories within a sample condition and between sample conditions. All models used were the most ‘brain-like’, as determined by the methodology in section “RNNs With STSP are More Brain-Like”. a) Cartoon of trial-averaged RNN trajectories corresponding to two different sample conditions for the fixed-synapse RNNs. a1) Average pairwise distance between trajectories on different sample conditions, for the fixed synapse network with tanh activation (FS-tanh). a2) The same plot as in a1, but for FS-relu. b1) The average distance between distracted and undistracted trajectories. Average taken over all sample conditions. Results shown for FS-tanh. b2) Same results as in b1, but for FS-relu.

The models with STSP did not achieve robust WM through persistent neural spiking, but instead relied on the STSP. We again examined trajectories using spikes rates from the STSP models (Figure 8). The neural distance between trajectories for different samples increased during sample presentation but then dropped back down near zero during the delay, especially longer delays. This mirrors the results from actual MUA activity in the PFC (see above). This stands in contrast to the fixed-synapse models (above), in which strong persistent spiking was unlike that observed in the brain. However, unlike the fixed-synapse models, in the STSP models we could use synaptic weights to measure trajectories and distance in *synaptic* state-space (i.e., decode the sample identity). This is analogous to using spike rates but instead we measure synaptic weights over time (as was done in Masse^33^). This revealed that trajectories for different samples in synaptic state-space remained elevated across the delay period (Figure 7), indicating sample information was being maintained by STSP even when spiking levels were low.

**Figure 8:**
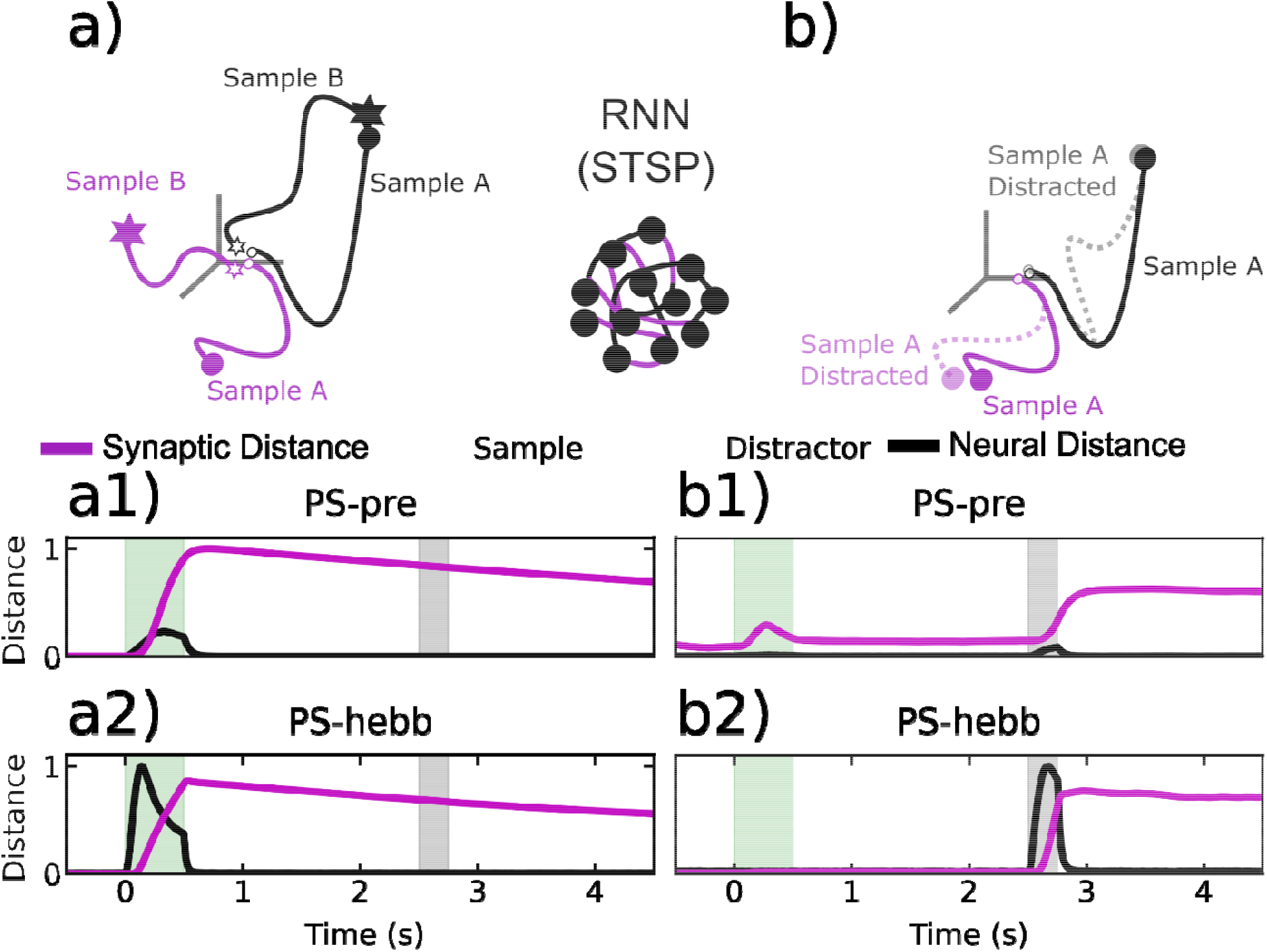
Distances between neural *and synaptic* trajectories within a sample condition and between sample conditions. Black lines correspond to neural trajectories, purple lines correspond to synaptic trajectories. All models used were the most ‘brain-like’, as determined by the methodology in section “RNNs With STSP are More Brain-Like”. a) Cartoon of trial-averaged neural and synaptic RNN trajectories corresponding to two different sample conditions for the plastic-synapse RNNs. a1) Average pairwise distance between neural and synaptic trajectories o*n different sample conditions, for* PS-pre. *a2) The same plot as in a1, but for* PS-hebb. *b1) The average distance between distracted and undistracted trajectories. Average taken over all sample conditions. Results shown for* PS-pre. *b2) Same results as in b1, but for* PS-hebb.

As we found in the experimentally recorded MUA, in the STSP models the spike rate trajectories between distractor and non-distractor trials increased during distractor presentation but then quickly decreased back to pre-distraction levels (Figure 8). By contrast, the distractor had a longer lasting effect in synaptic weight space for the STSP models (Figure 8). This is consistent with the distance plots of (Figure 8), which show the synaptic distance decreasing on a longer time-scale than the neurons. Since the synaptic distance decreased on a time-scale several times larger than the intrinsic time-constant of synapses, this indicates that the neurons and synapses interact in a way that increases their effective time constant. This could also potentially increase their susceptibility to perturbations. However, notably, we found that this was not the case. This increased time constant did not affect the ability of the STSP networks to perform the task. The STSP networks performed at a high level (above 90% accuracy) on both distractor and non-distractor trials.

## Discussion

We found that while RNNs with and without STSP showed robustness against distractors, the RNNs with STSP were more brain-like. We also found that RNNs with STSP were more structurally robust than RNNs without STSP. STSP models showed graceful degradation to synaptic loss. They were still able to function even after half of the synapses were ablated. By contrast, fixed-weight attractor dynamic models did not. They showed a severe decrease in function with as little as 10-20% synaptic loss. STSP models also showed more robust performance in response to noise. In the STSP models, decodability of spike rates decreased during the memory delay (especially longer delays). Importantly, the WMs *could* be decoded from the synaptic weights. This was in contrast to the RNNs without STSP, where spike-rate decodability remained high over the entire memory delay. This high spike-rate decodability did not match observations of actual spiking recorded from the PFC. In sum, STSP can not only maintain information over gaps of no spiking, it also adds functional advantages. And, notably, adding STSP to RNNs makes them exhibit brain-like behavior.

Some forms of STSP worked better than others. We found that purely Hebbian STSP models were difficult to train, in agreement with previous studies^34^. By contrast, an anti-Hebbian STSP model (which reduced synaptic weights when spiking was too correlated) was successful and did not require any external weight clipping. We and others have found that anti-Hebbian STSP plays a crucial role in network stability and function^31,35^. And like others, we found that the synaptic weights in the *PS-pre* model had to be limited to a certain range to maintain stability and trainability^19,33^. This assumption is biologically plausible. Biological synapses are limited in the amount of resources that can be recruited by a presynaptic spike^19,36^.

The STSP models are in line with a variety of studies suggesting that cognition is more complex than steady attractor states. For example, sustained attention was long-thought to depend on attractor-like steady-state spiking. However, like studies of WM, this may have been an artifact of averaging spiking across trials^11,14^. Examination of sustained attention in real time (on single trials) has shown that, behaviorally, attention waxes and wanes rhythmically at theta^37^. There is a corresponding waxing and waning of spiking synchronized to LFP theta oscillations. Likewise, neural correlates of WM show sparse bursts of spiking linked to oscillatory dynamics when examined in real time^14,15^.

Whether WM relies on persistent attractor dynamics (i.e., persistent spiking) alone or sparse attractor dynamics with STSP is of current interest^9–11,38,39^. A recent computational study has provided insight. Masse et al trained a WM model with STSP ^33,40^. They found that there was sparse spiking when WMs were being simply maintained and more spiking when WMs were being used and manipulated. This is consistent with other STSP-based WM models. For example, Lundqvist et al^14^, found that spiking increases when WMs are being read out for use. This makes sense because the brain cannot read information from the synapses directly. Spikes are needed to “ping” the network to read information from the synaptic weights. Thus, models with STSP can work for both modes: Sparse spiking when WMs are being maintained and more spiking when WMs are being used. It is also worth noting that the brain may use other mechanisms for performing WM tasks beyond STSP and elevated spiking, for example rotational dynamics ^41^.

It is also worth noting that studies that report higher levels of memory-delay spiking tend to be those that used spatial delayed response tasks. Spatial delayed response may involve more motor inhibition than WM per se^15^. The animal knows the forthcoming behavioral response and is inhibiting it while waiting for a “go” signal. It is like revving the engine while keeping your foot on the brake. By contrast, WM tasks like delayed match-to-sample (i.e., the task used in this study), the behavioral response is not known until after the memory delay. They involve holding information for further computation, not inhibiting a behavioral response. Tasks like delayed match-to-sample are well known to produce lower levels of memory delay spiking^14,15,33,42^ and thus require more than attractor states alone.

The robustness and stability added by STSP is critical not just to WM but to top-down control in general. A control system which is overly sensitive to perturbations would quickly propagate these disturbances to the rest of the brain, leading to errant behavior. Moreover, robustness and stability are intimately tied to modularity^43–45^. Stable networks can be linked with other stable networks in a way that preserves stability. Without that property, the whole network must be retrained when new modules are added to subserve complex or composite behavior. Our results suggest that STSP has a dual role. It can maintain information while simultaneously ensuring robustness in an energy-efficient manner^9^. We speculate that stability could be a key component to understanding the way specialized modules in the brain dynamically cooperate to form cohesive perceptions, plans, and actions.

## Methods

### Subject and Task

The non-human primate subject used in our experiments was a male rhesus macaque monkey (*Macaca mulatta*), aged 17. All procedures followed the guidelines of the Massachusetts Institute of Technology Committee on Animal Care and the National Institutes of Health. As described in the main text, Utah arrays were implanted bilaterally in dorsolateral PFC (dlPFC) and ventrolateral PFC (vlPFC). Animal performed a distracted delayed match-to-sample task. Each trial began with visual fixation on the middle of the screen for 0.5s. Fixation was maintained throughout the trial until the behavioral response. The delay length was parametrically varied from 1 – 4 s in five logarithmic steps, randomly chosen each trial. At mid-delay a neutral distractor (1 of 2 possible objects never used as samples) was presented randomly on 50% of trials. During the multi-choice test the monkey was allowed to freely saccade between all objects on the screen. The final choice was indicated by fixating on it for at least one second.

### Data Analysis Methods

#### Distances Between Neural Trajectories

All neural spiking data was first smoothed using a 10ms Gaussian kernel. We refer to the smoothed spiking data as the firing rate. To quantify the difference in population activity between different conditions, we computed the distance between neural trajectories. We define a neural trajectory as a vector of firing rates for *N* recorded neurons evolving in time.

For the distractor vs non-distractor comparison, denote 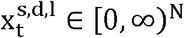 the neural trajectory at time *t* on trial l ∈ {1,…, L} for stimulus identity s ∈ {1,…, S} and where d ∈ {0,1} denotes the presence or absence of a distractor in the delay phase. We first computed trial averages as

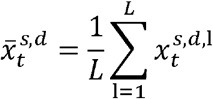

Then for each *s* we computed the distance between distracted and non-distracted trajectories, using the Euclidean norm, and took the average over stimuli:

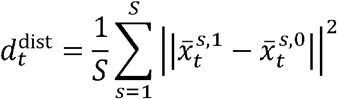

For the comparison between trajectories of different stimulus identities, we used only non-distractor trials (*d* = 0). We first averaged over trials as above for each stimulus to get

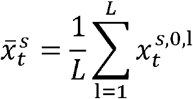

and calculated the mean distance between each pair of stimuli as:

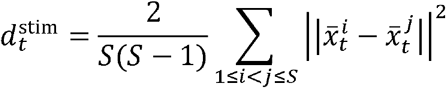

#### Confidence Intervals for Trajectory Distances

As our final measurement we calculated the mean over six experimental sessions for both 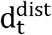 and 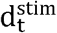. To quantify uncertainty that captures both the session and trial variability, we used a non-parametric multi-level bootstrap. Briefly, for b ∈ 1, …, B with *B* = 1000, we first sampled with replacement over sessions (that is, we randomly selected a session with replacement *E* times, where *E* is the total number sessions in the dataset - in our case *E* = 6), and then for each resampled session, we resampled trials with replacement in the trial-average steps described above, yielding the bootstrap samples 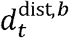 and 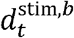. We took the 2.5^th^ and 97.5^th^ percentiles across *b* as our lower and upper values for the 95% confidence interval.

#### Sample Decoding

For sample decoding experiments, we first smoothed trials (as above, with 10ms Gaussian kernel), and then took average rates per neuron in 50ms time bins and attempted to decode sample type from the population activity at each time point. For our classifier, we used a linear Support Vector Machine and standard regularization (C=1 in SciKit Learn). For each session we used a 10-fold cross-validation to calculate decoding accuracy. (That is, we held out 10% of trials as a test set, and trained on the remaining 90% of the data. We did this for 10 non-overlapping test sets, and then took the average test accuracy). We performed a decoding analysis on each session separately, and then took the average and standard error across sessions for our result.

#### Change from Baseline Decoding

We smoothed trials and took average rates as in the sample decoding above. Then, for a given sample type, we took the population activity from 400-350ms before the stimulus (or distractor) and attempted to decode subsequent population activity against this baseline activity. We performed 10-fold cross-validation as above for calculating test accuracy, and calculated the average performance over samples for each session. Plots show the average and standard error across sessions.

#### Sensitivity to Smoothing and Bin Size

Throughout our data analysis we used a 10ms Gaussian kernel to produce firing rates, and in the decoding analysis, we used the mean firing rate in 50ms time bins. To ensure our trajectory distance results did not depend specifically on the choice of Gaussian kernel, we examined trajectory distances using 5, 10, and 15ms Gaussian kernels (Figure S6). For the distance between trajectories with different sample identities, we took the mean distance within the sample presentation period (0, 0.5*s*) and the 100ms preceding test time (*t*_*test*_ − 0.1*s, t*_*test*_), for each delay and each session, and then compared the sample distances and test distances. Similarly, for distances between distracted and non-distracted trajectories, we took the mean distance within the distractor presentation (*t*_*dist*_, *t*_*dist*_ + 0.25*s*), and the (*t*_*test*_ − 0.1*s, t*_*test*_), again for each delay and each session. To ensure our decoding results did not depend on the specific choice of Gaussian kernel and length of time bin, we again used 5, 10, and 15ms Gaussian kernels and additionally used 20ms, 50ms, and 100ms time bins (Figure S7). We then looked at sample decoding accuracy in the non-distracted trials, for each combination of Gaussian kernel and time bin, again taking a mean for the sample presentation (0, 0.5*s*), and a mean for the pre-test period (*t*_*test*_ − 0.1*s, t*_*test*_), for each delay and each session. The Wilcoxon signed-rank test was used for comparisons between time epochs (*e*.*g*. Sample vs Pre-Test), with p-value corrections for multiple comparisons made using the Benjamini-Hochberg procedure (with *α* = 0.05).

### Artificial Neural Network Modelling

#### Fixed-Synapse (FS)

For the fixed-synapse ANNs we used a ‘vanilla’ recurrent neural network. Each neuron had an activation *x*_*i*_ which we collected into a vector *x* ∈ *R*^*n*^ such that (*x*)_*i*_ = *x*_*i*_ This vector was the *state* of the neural network at a given time *t*. The state evolves in time according to the following differential equation:

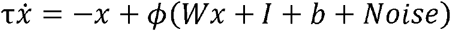

Here *τ* is the time-constant of the network, which we set to 100 milliseconds. The function *ϕ*(·) is a nonlinearity (e.g tanh, softplus, etc). The matrix *W* is the recurrent weight matrix of the network, and determines how the different neurons interact with one another. The vector *I* ∈ *R*^*n*^ is the exogenous input into the recurrent neural network, and can vary with *t*. For our purposes *I* will represent sensory information related to the task at hand. The vector *b* ∈ *R*^*n*^ is a fixed bias term. The noise will be defined shortly. To integrate the neural dynamics, we use the standard Euler approximation scheme with fixed step-size of *dt* = 15 milliseconds. Defining the ratio of *dt* to *τ* as 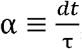, the approximated dynamics are:

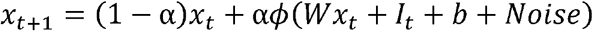

Here 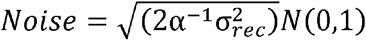 where *σ*_*rec*_ = 0.05 denotes the *process noise* and *N*(0,1) denotes a draw from the standard normal distribution. For the *FS-tanh* networks, we use *ϕ*(·) = tanh(·). For the *FS-relu* networks, we use *ϕ*(·) = ReLU(·) = max(0,·). Finally, we used an affine readout for the network to produce the desired output for the task:

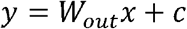

With the exception of *W*, all the parameters of the fixed-synapse networks were initialized by drawing each element randomly from a gaussian distribution with mean zero and standard deviation 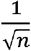. For the recurrent weight matrix *W* the elements were drawn from a gaussian distribution with mean zero, but standard deviation 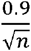. We found that this initialization achieved faster training. At the beginning of each trial, the network state was initialized at the origin (i.e *x* = 0).

#### Plastic Synapse

##### PS-pre

This ANN was an implementation of the model originally introduced by Mongillo et al ^19^ and then subsequently trained using deep learning techniques by Masse et al ^33^. Many of the details are similar to or exactly the same as in Masse et al. We go through them here for completeness. As in the fixed-synapse model, we used Euler integration to approximate the dynamics:

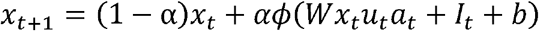

Aside from the addition of two new variables *u*_*t*_ and *a*_*t*_ (which we’ll define shortly) this is the same model as the fixed-synapse model. This network consisted of 80% excitatory neurons and 20% inhibitory neurons. To keep these populations separated throughout training, we decomposed *W* as 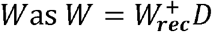 where 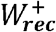 is a matrix with non-negative entries and *D* is a diagonal matrix whose *ii*^*th*^ entry is either 1 or −1, depending on whether neuron *i* is an excitatory or inhibitory neuron. To each synapse we associated a value *a*, for the fraction of available neurotransmitter and *u*, the amount of utilized neurotransmitter. These two variables evolved according to:

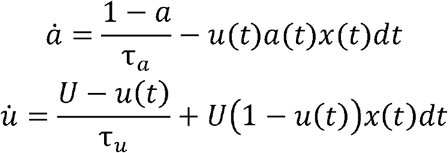

here *x*(*t*) is the presynaptic activity at time *t, τ*_*a*_ is the time constant of available neurotransmitter recovery, *τ*_*u*_ is the time constant of neurotransmitter utilization recovery. Half the synapses in the network were facilitating, the other half was depressing. For the facilitating synapses, *τ_a_* = 200 *ms*, *τ_u_* = 1500 *ms* and *U* = 0.15. For the depressing synapses, *τ_a_* = 1500 *ms*, *τ_u_* = 200 *ms* and *U* = 0.45. Aside from 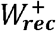, *t*he parameter initialization for *PS-pre* was the same as the fixed synapse networks. For *W*, we found that the following initialization worked best in terms of achieving the fastest training time:

1. Draw each element of 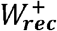 from the log-normal distribution, with the underlying normal distribution having mean 0 and standard deviation 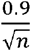.
2. Compute the largest singular value of 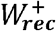 (call it *γ*) and divide every element of 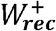 by 10*γ*.

To ensure that 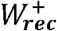 stayed non-negative throughout training, we always ran each element through a rectified linear function at the beginning of the trial. For each trial, the neural state vector *x* was initialized at the origin. The facilitation variables *a* were all initialized at *a* = 1. The utilization variables were all initialized at *u* = 0. Throughout the trial, both facilitation and utilization were clamped between 0 and 1. The readout from the network was affine, as in the fixed-synapse networks.

##### PS-hebb

This excitatory anti-Hebbian / inhibitory Hebbian ANN was originally introduced in Kozachkov et al ^31^. As before, we use a Euler integration scheme to approximate the dynamics:

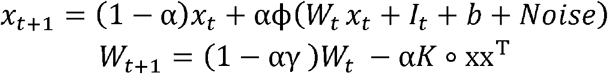

Here ϕ(·) = ·. That is, ϕ is just the identity function. The parameter γ controls the degree of weight decay, and is set to 1/200*ms*^−1^ unless otherwise noted. The matrix *K* is positive and positive-definite, and the symbol ∘ denotes element-wise matrix multiplication (i.e Hadamard product). During training we ensure positiveness and positive-definiteness of by parameterizing it as:

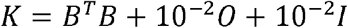

Where 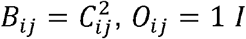 is the identity matrix. Since *B*^*T*^*B* is positive semi-definite and non-negative, the matrix *O* ensures that *K* is strictly positive, while the identity matrix ensure that *K* is strictly positive-definite. The training is done on the matrix *C*. The output from the network is affine, as in all the other networks. All the parameters are initialized the same way as the fixed-synapse networks, except for *C*. The elements of this matrix are drawn from a uniform distribution between −0.5 and 0.5. At the beginning of each trial, the neural state vector *x* and the weights *W* are initialized at the origin.

For all models, we tested how the numerical integration step-size impacted performance. We used the hyperparameters corresponding to the most brain-like models. For each of the models, we tested a smaller step-size (10ms) and a larger step-size (30ms). All the models expect *PS-hebb* performed the same. *PS-hebb* was sensitive to the larger step-size, but insensitive to the smaller step-size. This makes sense because the stability guarantees for *PS-hebb* were proven in continuous-time. Smaller step-sizes provide more accurate approximations to the precise continuous-time solution.

We found that the initialization of the parameters was important for all models. For the fixed-synapse models, initializing the weight matrix with a slightly sub-critical matrix yielded a good tradeoff between initial stability and expressivity. For the *PS-pre* model, using the log-normal initialization provided by Masse worked well. When we experimented with different initializations, training either failed or was slower. The same was true of the *PS-hebb* model—different initialization schemes led to different training profiles in terms of model performance and training time.

#### Task Structure and Training Details for Artificial Neural Networks

The basic task structure for the ANNs was the same as for the non-human primate. Each stimuli was encoded in a one-hot vector. There was a total of 10 images (8 sample images, 2 distractor). The number of input neurons going into the ANNs was 11 = 10 + 1: 10 input neurons for the stimuli and 1 for a fixation signal. These input stimuli were collected into a vector *m*_*t*_, which changed with time throughout the trial. This vector was fed into the ANNs via a trainable input weight matrix *I*_*t*_ = *W*_*in*_ *m*_*t*_. There were 11 = 8 + 2 + 1 output neurons leaving the ANNs, 8 for each sample image, 2 for the distractors and one as a fixation neuron. These neurons were collected into the vector *y*_*desired*_, which defined the desired output of the ANNs.

At *t* = 0 the ANN has to fixate by having its fixation output neuron hold a value of ‘1’. The fixation is left on for 1000ms. At the end of the fixation period a randomly chosen sample image is presented for 500ms. Following the sample presentation period, the delay period begins. The delay length is randomly chosen from 1.0 s, 1.41s, 2.0 s, 2.83 s and 4s.

On 50% of the trials a distractor is presented in the middle of the delay for 250ms. Following the end of the delay period, the test period begins. During the test period the original sample image is presented as well as an off-target image for 500ms. At this point the RNN must do two things: stop fixating (i.e make the output fixation neuron output a value of ‘0’) and indicate the correct choice by making the output neuron corresponding to the sample image hold a value of ‘1’. We use a mean-squared loss to quantify network performance. For each trial we take the network output for a period of 500ms after the test period ends and compute the loss:

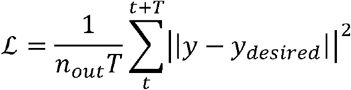

We compute this loss for each trial in the batch, and then average over the batch to get a total loss per batch. Our training and test sets consisted of 2^14^ trials each. We used a batch size of 256 to train and test. To train the networks we use the Adam optimizer with Pytorch standard parameters, a weight-decay value of 10^−4^ for all models unless otherwise specified. For the FS models we use a learning rate of 10^−3^. For *PS-pre* we use a learning rate of 0.02. For *PS-hebb* we use a learning rate of 0.01.

As described in the main text, we concluded a hyperparameter search to assess the generality of our claims. For reasons we will describe shortly, the *PS-hebb* model was treated slightly differently than the other three. For *FS-relu, FS-tanh* and *PS-pre*, we looped over:

- Hidden size = (100,200,300,400,500,600,700,800,900,1000)
- Activity regularization = (0.0, 0.5, 1,1.5,2,2.5)
- Parameter regularization (weight decay) = (1e-6, 1e-5,1e-4,1e-3,1e-2)

The activity regularization, we computed the squared sum of all the neuron activations in a batch, and divided by the number of neurons and the number of time points. This quantity was then scaled by the regularization strength parameter which was varied during the hyperparameter sweep. This scaled quantity was then added to the overall loss function. For the *PS-hebb* model, we only varied the hidden size from 10 neurons to 100 neurons in steps of 10, and we additionally varied the γ parameter (synaptic decay strength) between (.001,.01,0.1,1). The reason we could not increase the size of the model to 1000 is due to GPU memory issues. The number of variables in *PS-hebb* model scales quadratically with the number of neurons (because each synapse is also a variable).

To test network performance, we compute the ‘decision’ of the network as the index of the output neuron which had the highest value. We compute the average accuracy of the network by the following procedure:

1. In the 500ms following the end of the test period, we compare the network output to the desired output for each time-point. We add up the number of times the network makes the right decision in this period, and then divide over the number of time-points to get a single-trial accuracy.
2. We compute this single-trial accuracy for all trials in the test set, and report the final network accuracy on the task as the average single-trial accuracy, taken over the whole test set.

## Supplementary Figures

For all supplementary figures, green indicator box denotes sample period, gray distractor period, blue test period. Dotted lines indicate chance accuracy values

**Figure S 1:**
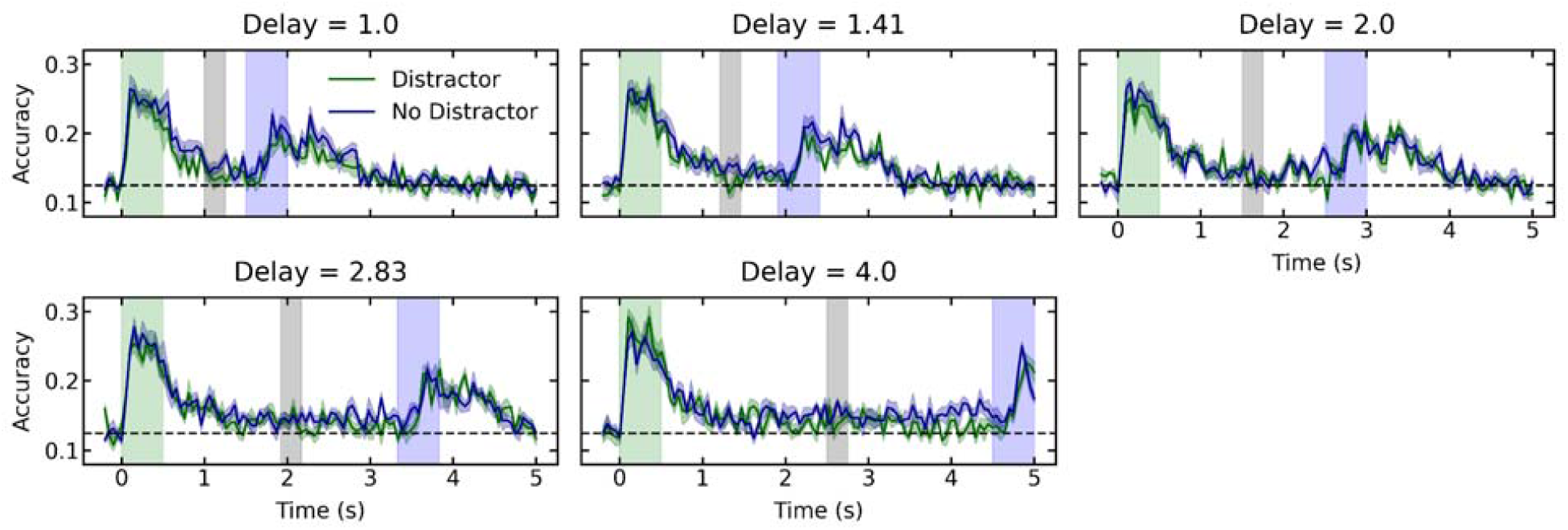
Decoding sample identity from neural spiking, Classifier was a linear support vector machine. We evaluated performance on held-out data. Sample decoding accuracy drops to approximately chance during the delay

**Figure S 2:**
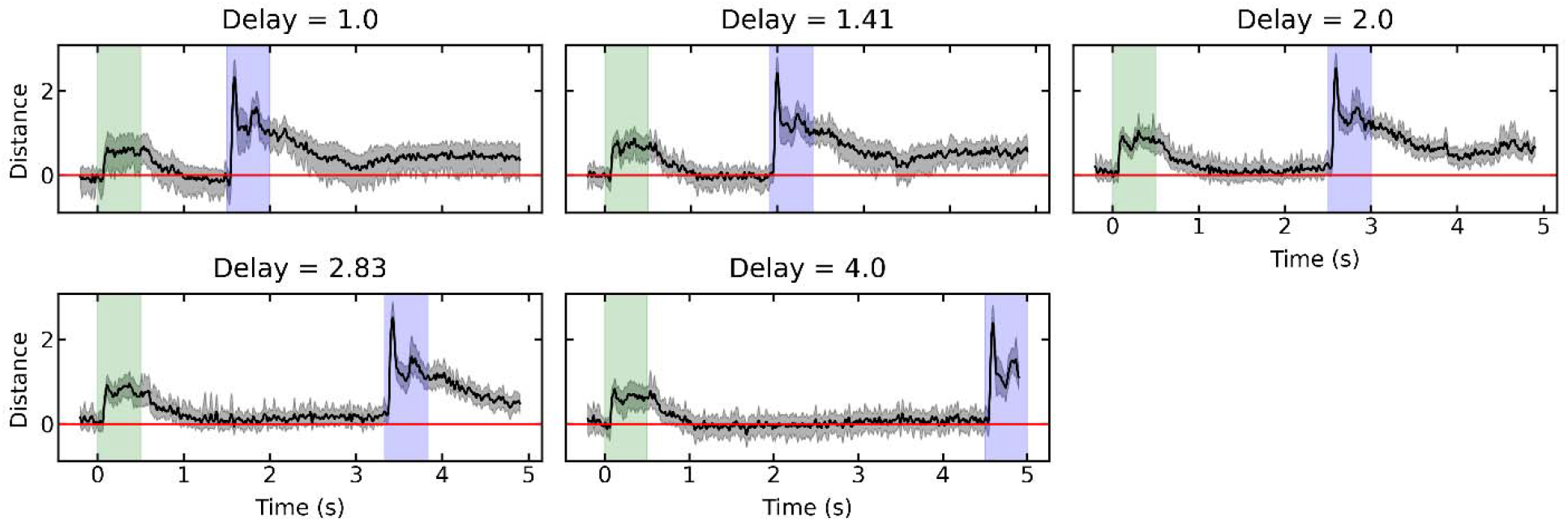
Average distance between pairs of trajectories elicited by different samples.

**Figure S 3:**
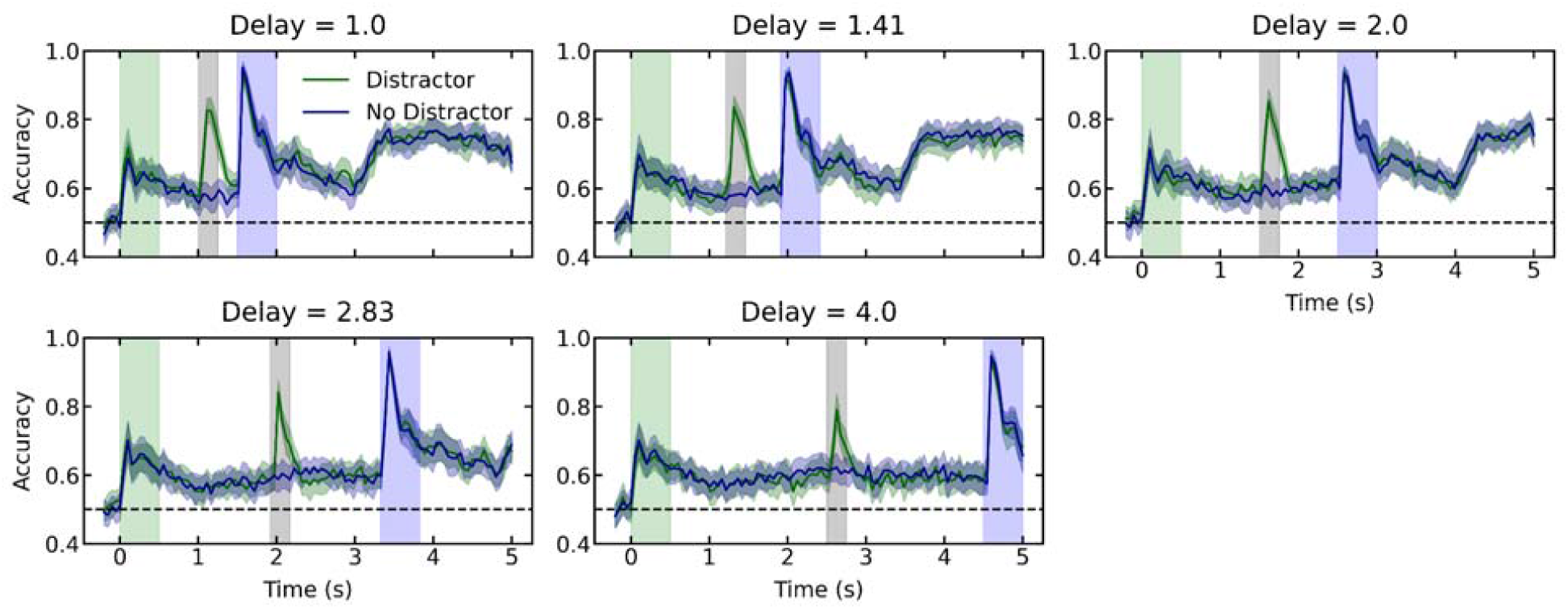
Decoding pre vs post sample presentation from neural spiking. Spiking contains information about pre vs. post sample presentation through the delay. This indicates that neural spiking does not return to its pre-sample firing pattern following sample presentation.

**Figure S 4:**
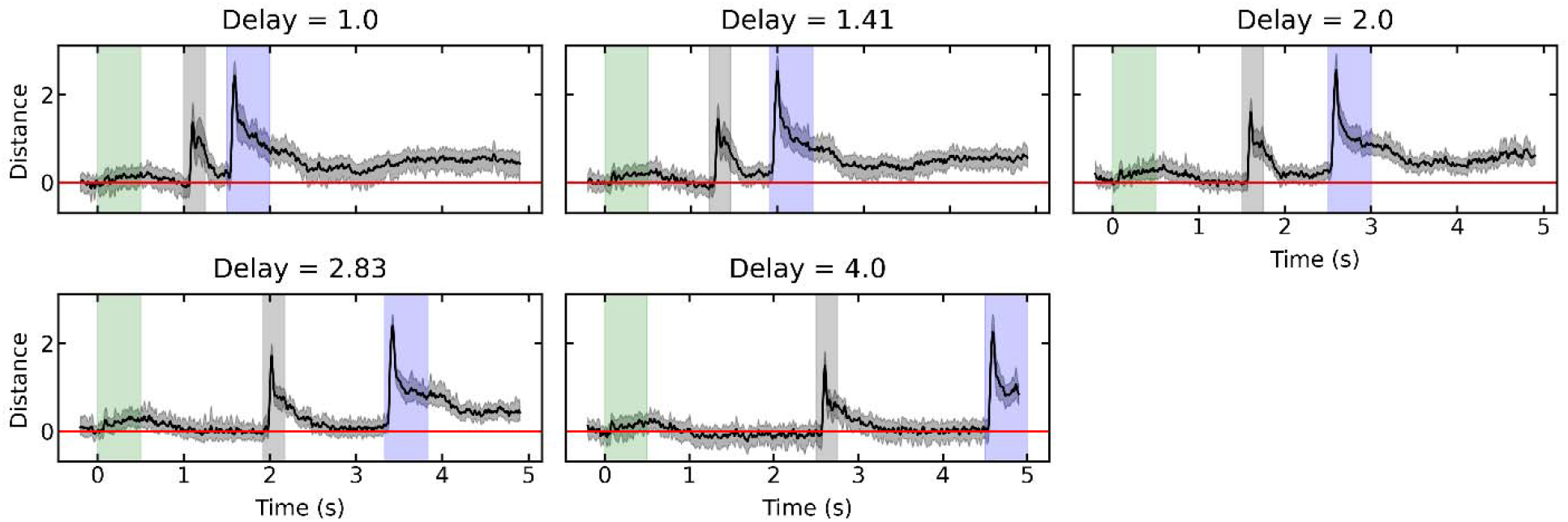
Average distance between distracted and non-distracted trajectories corresponding to the same sample item.

**Figure S 5:**
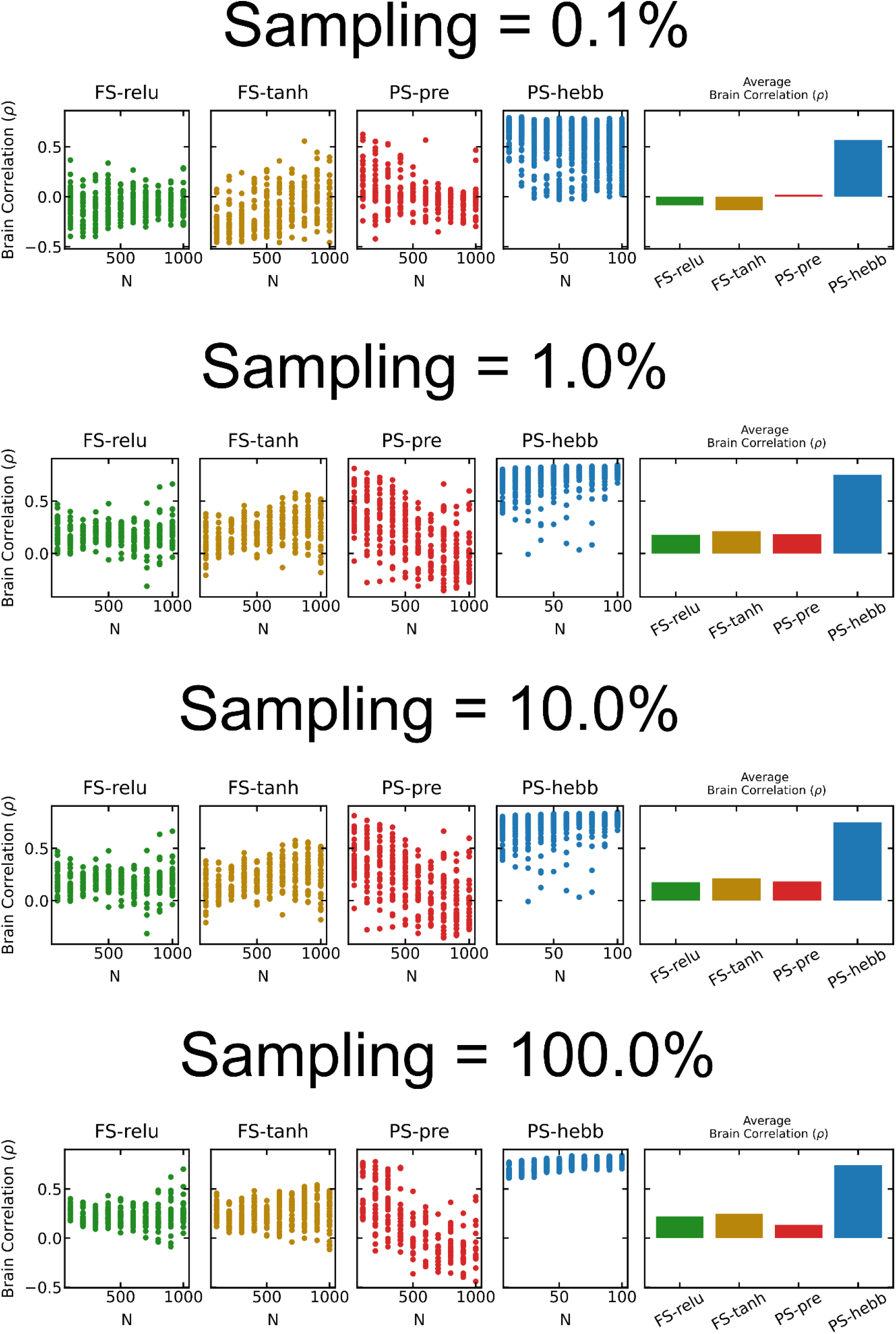
The effect of subsampling the RNNs on the brain-correlation index used. The qualitative result stays the same (PS-hebb is more brain-like on average), but the particular correlation values change.

**Figure S 6:**
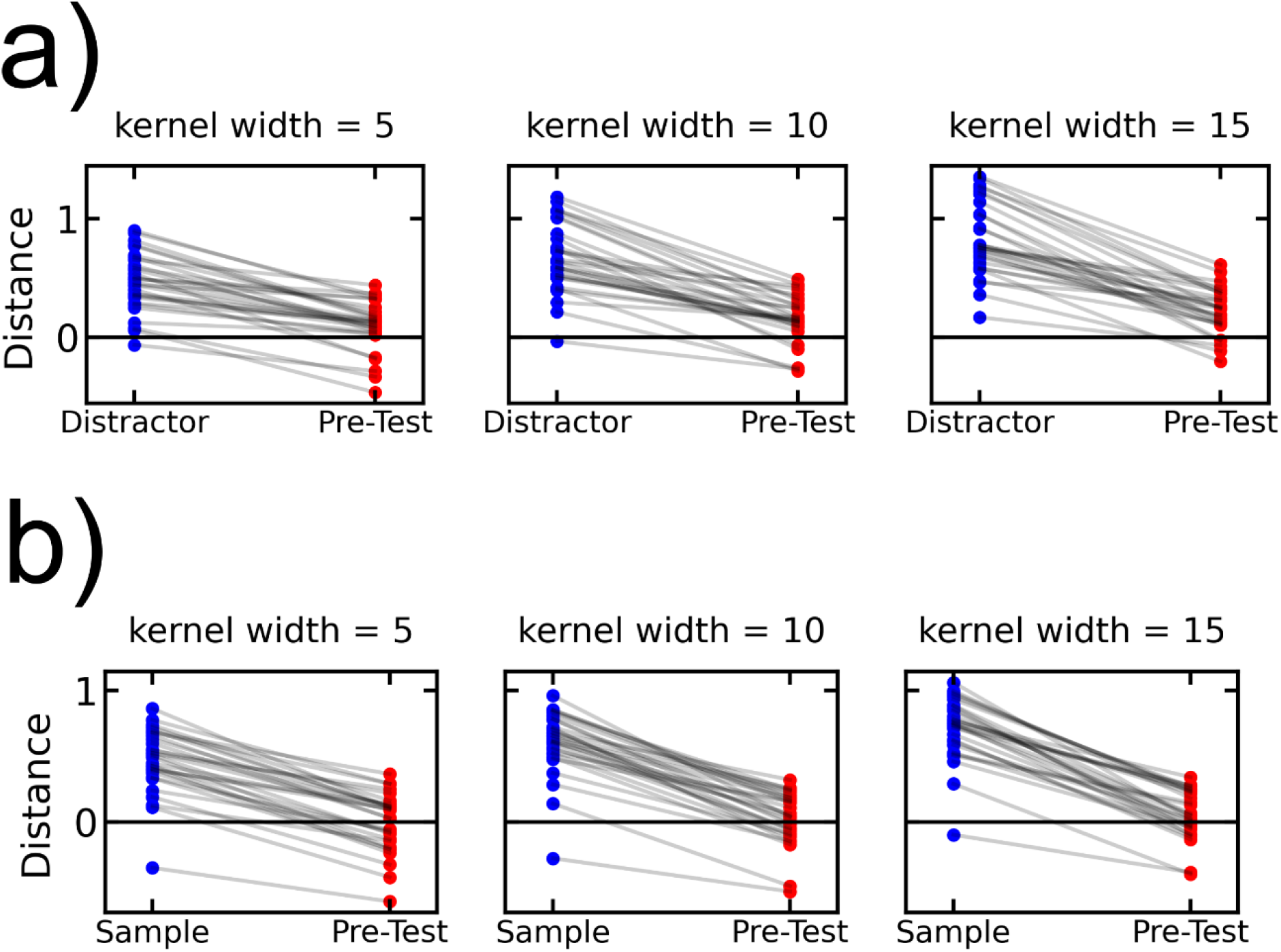
Effect of using different smoothing parameters on trajectory distances. In all plots, each pair of dots correspond to averages during two time epochs for a particular session and delay time. In both cases the Pre-Test epoch used was 100ms preceding test time. All comparisons were statistically significant (p < 1e-3, Wilcoxon signed-rank test). a) Average distance between neural trajectories for different sample IDs for different smoothing parameters. Comparison is between Sample epoch (blue) and Pre-Test epoch (red). b) Average distance between neural trajectories for distracted and non-distracted trials for different smoothing parameters. Comparison is between Distractor epoch (blue) and Pre-Test epoch (red).

**Figure S 7:**
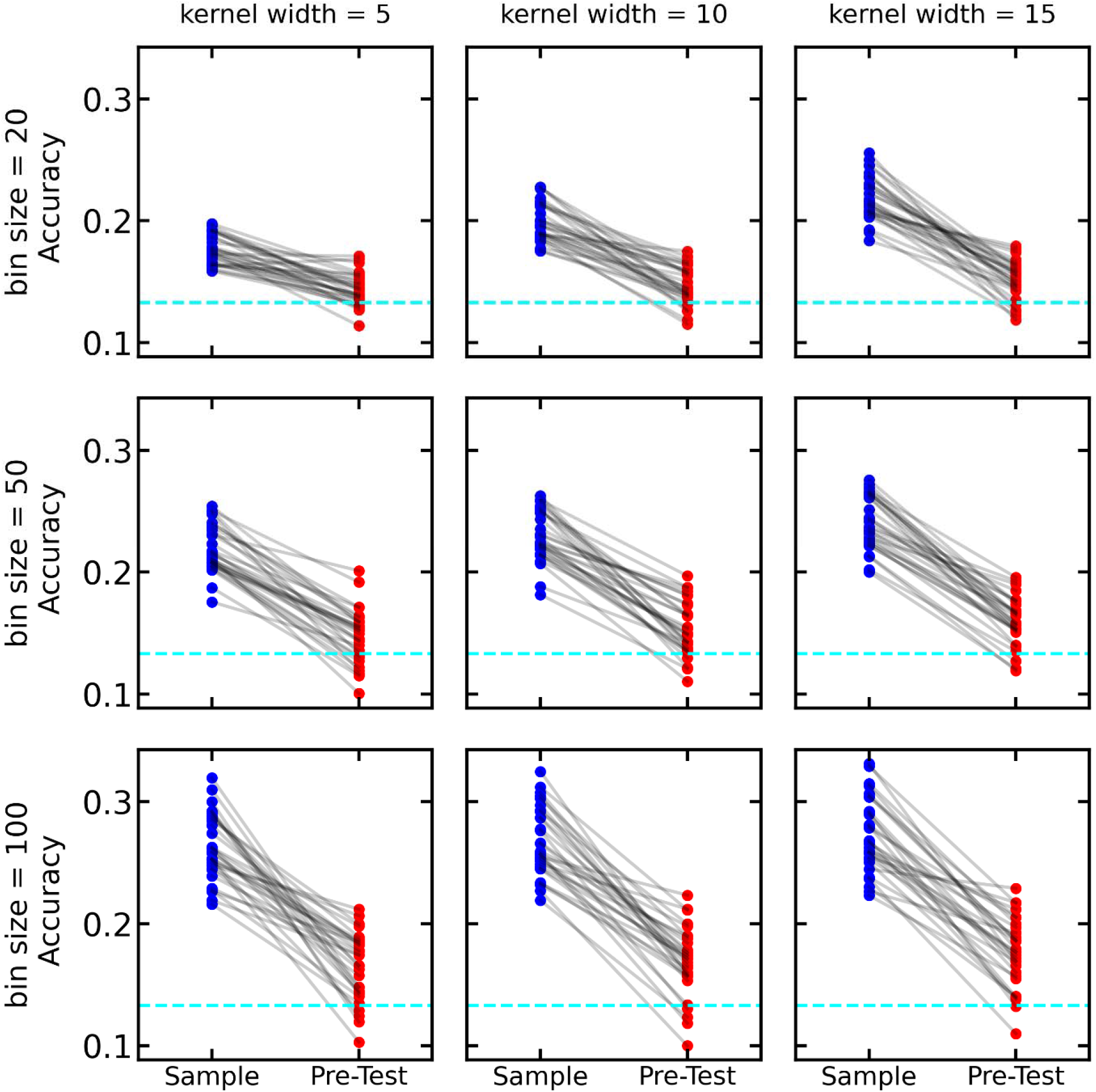
Average decoding accuracy on non-distractor trials for different smoothing parameters and different time bin sizes. Comparison is between Sample epoch (blue) and Pre-Test epoch (red). In all plots, each pair of dots correspond to averages during two time epochs for a particular session and delay time. The Pre-Test epoch used was 100ms preceding test time. All comparisons were statistically significant (p < 1e-3, Wilcoxon signed-rank test).

**Figure S 8:**
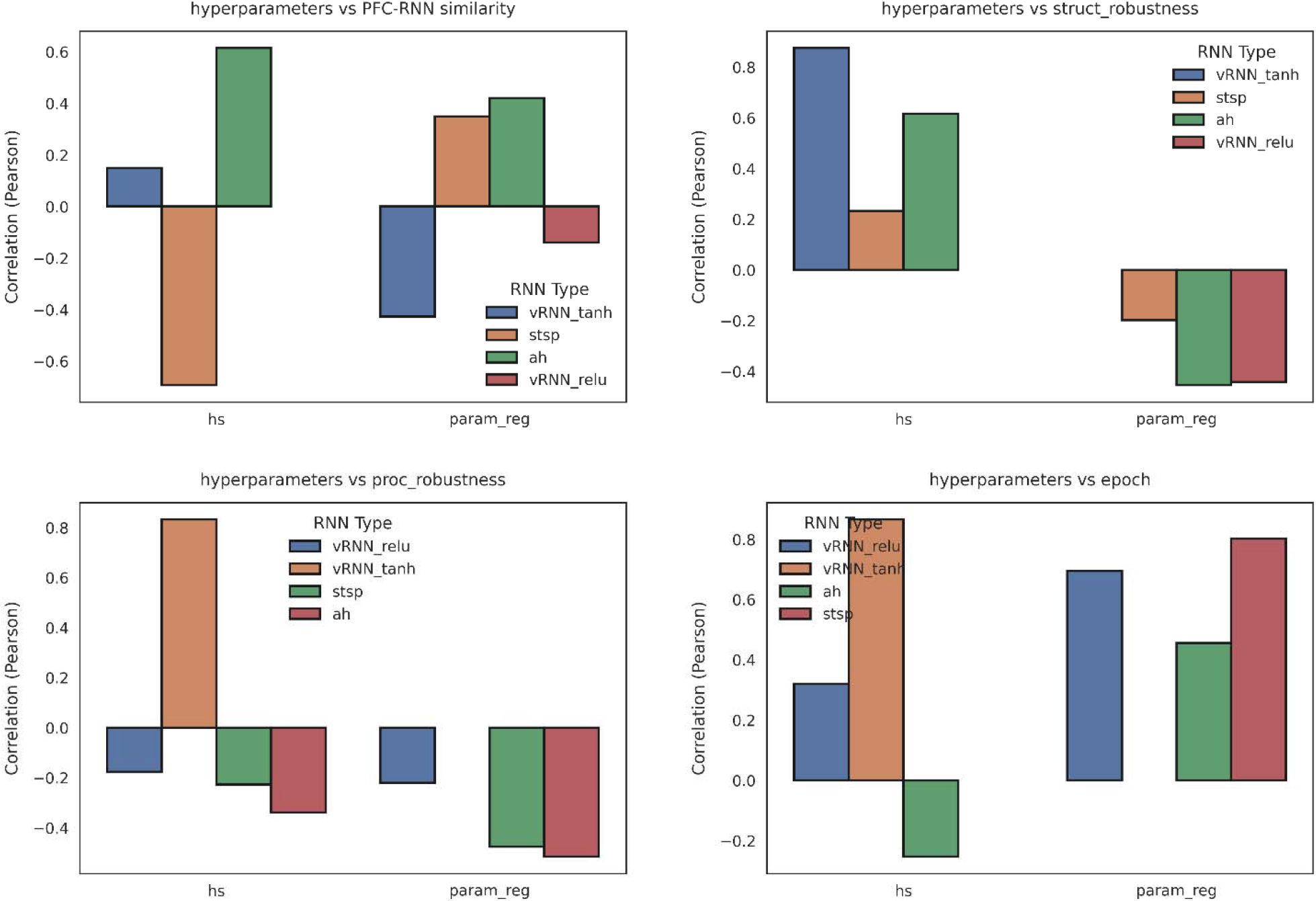
We measured the impact of each hyperparameter (e.g hidden size) on dependent variables of interested (e.g structural robustness) by fitting a linear trend. In this figure we only report significant (p < .005) relationships.

